# Long-read genome assembly of the Japanese parasitic wasp *Copidosoma floridanum* (Hymenoptera: Encyrtidae)

**DOI:** 10.1101/2023.09.24.559078

**Authors:** Kouhei Toga, Takuma Sakamoto, Miyuki Kanda, Keita Tamura, Keisuke Okuhara, Hiroko Tabunoki, Hidemasa Bono

## Abstract

*Copidosoma floridanum* is a cosmopolitan species and an egg-larval parasitoid of the Plusiine moth. *C. floridanum* has a unique development mode called polyembryony, in which thousands of genetically identical embryos are produced from a single egg. Some embryos develop into sterile soldier larvae, and their developmental patterns differ between the US and Japanese *C. floridanum* strains. Genome sequencing can accelerate our understanding of the molecular bases underlying polyembryony, including the production of soldier castes. However, only the genome sequence of the US strain has been reported. In the present study, we determined the genome sequence of the Japanese strain using Pacific Biosciences high-fidelity reads and generating a highly contiguous assembly (552.7 Mb, N50: 17.9 Mb). Gene prediction and annotation identified 13,886 transcripts derived from 10,786 gene models. We searched the genomic differences between US and Japanese strains. Among gene models predicted in this study, 100 gene loci in the Japanese strain had extremely different gene structure from those in the US strain. This was accomplished through the functional annotation (GGSEARCH) and long-read sequencing. Genomic differences between strains were also reflected to amino acid sequences of *vasa* that plays a central role in caste determination in this species. The genome assemblies constructed in this study will facilitate the genomic comparisons between Japanese and US strains, leading to our understanding of detail genomic regions responsible for the ecological and physiological characters of *C. floridanum*.

## Introduction

*Copidosoma floridanum* (Hymenoptera: Encyrtidae) is a worldwide distributed egg-larval parasitoid of the Plusiinae moth (Lepidoptera, Noctuidae, Plusiinae) (Guerrieri and Noyes 2005). This species reproduces by polyembryony, producing >2,000 genetically identical individuals from a single egg (Ode and Strand 1995). Polyembryony has evolved independently in the four Hymenoptera families (Encyrtidae, Platygastridae, Braconidae, and Dryinidae); however, encyrtid wasps have the most extreme form regarding clone size (Segoli et al. 2010). Embryos develop into larvae that feed on the host tissues until pupation. In some encyrtid wasps, the embryos develop into two types of larvae: reproductive larvae and the sterile soldier larvae. Sterile soldier larvae possesses large mandibles, and contribute to the defense of their clonal siblings by attacking against a competitor in the host (Cruz 1981; Strand 2003; Iwabuchi 2019). The phenomenon of solitary individuals forming cooperative groups with a reproductive division of labor can be considered as one of the major evolutionary events that brought complexity to life on Earth (Szathmáry and Smith 1995; Whyte 2021).

The process and molecular basis of soldier development in *C. floridanum* have been described and investigated mainly in the US and Japanese strains (Ode and Strand 1995; Smith et al. 2017; Sakamoto et al. 2020). Like other hymenopteran insects (Cook 1993), *C. floridanum* is haplodiploid species in which the females develop from the fertilized eggs and the males develop from the unfertilized egg (Iwabuchi 2019). Soldier larvae develop from both male and female embryo, and the development and behavioral patterns of male soldiers differed between intraspecific strains. In the US strain, male soldiers appear in the penultimate larval stage of the host (Grbic et al. 1997), whereas in the Japanese strain, they appear in the first- or second-instar larvae of the host (Yamamoto et al. 2007). Male soldiers were functional in the Japanese strain but are not as aggressive as the female soldiers (Uka et al. 2013). Male soldiers in the US strain are nonaggressive (Giron et al. 2007). *vasa*, encoding a DEAD-box RNA helicase protein (Hay et al. 1988; Lasko and Ashburner 1988), has been shown to play a central role in caste determination of *C. floridanum* (Donnell et al. 2004; Vladimir Zhurov et al. 2004). The *vasa* gene was originally identified in *Drosophila melanogaster* as a maternal-effect factor for germ-cell specification and development of abdominal segments (Schupbach and Wieschaus 1986), and they have been found in invertebrates and vertebrates (Raz 2000). *vasa* has been compared between the Japanese and the US strains of *C. floridanum*, and a 10 amino acid sequence difference in the N-terminal region was observed (Ohno et al. 2019).

Each strain should optimize its behavior and developmental patterns for its habitat. Genome or transcriptome analyses should accelerate our understanding of the molecular basis underlying the intraspecific differences of parasitic ability including polyembryony and caste system (Sakamoto et al. 2020). The genome of the US strain has been sequenced using short-read sequencing (i5K Consortium 2013), while that of the Japanese strain has not. Long-read sequencing offers several advantages over short-read sequencing (Pollard et al. 2018). High-quality (high contiguity of the assembly and completeness of gene sets) insect genomes have recently been achieved using long-read sequencing (Hotaling et al. 2021). Single-molecule real-time (SMRT) sequencing developed by Pacific Biosciences (PacBio) employ the DNA polymerase reaction from a single circular DNA template (Eid et al. 2009), generating highly accurate long high-fidelity reads (HiFi reads, >10 kb, >99%) (Rhoads and Au 2015; Ardui et al. 2018; Wenger et al. 2019). HiFi reads are more suitable than other long- or short-read methods for identifying genes containing repetitive sequences (Kawahara et al. 2022; Hotaling et al. 2023).

Comparisons among intraspecific genomes should be useful in understanding of the various parasitic ability in *C. floridanum*. In this study, we aimed to construct a highly contiguous genome assembly of the Japanese *C. floridanum* using PacBio long-read sequencing. We performed the quality check, the gene prediction and functional annotation for the *de novo* assembly. Gene annotation and amino acid sequences of *vasa* were compared to clarify the genomic differences between Japanese and U.S. strains.

## Materials and Methods

### DNA extraction and genome sequencing

Genomic DNA was extracted from adult *C. floridanum* males using the Blood & Cell Culture DNA Maxi Kit (Qiagen Co. Ltd., Valencia, CA, USA). The genetic system of *C. floridanum* and other hymenopteran insects is haplodiploid; females develop from fertilized eggs (diploid), and males develop from unfertilized eggs (haploid). In this study, the male adults were used. The male adults are clones born from a single egg and are genetically nearly identical. The sequence library was prepared using the SMRTbell Express Template Preparation Kit 2.0 or SMRTbell Prep Kit 3.0 (Pacific Biosciences, Menlo Park, CA, USA). Libraries were sequenced using the Sequel II or IIe (Pacific Biosciences).

### *De novo* assembly and quality assessment

Subreads and HiFi reads generated from the PacBio sequencers were assembled using Flye v 2.9 (Kolmogorov et al. 2019) and Hifiasm v 0.16.1 using the ‘-primary’ option (Cheng et al. 2021), respectively. In the PacBio long-read system, polymerase reads are yielded by sequencing circularized DNA. Then, the adaptors are removed from the polymerase reads, offering the subreads (refer to https://www.pacb.com/technology/hifi-sequencing/). HiFi reads are defined as the consensus sequence called from the subreads. Hifiasm v 0.16.1 can produce the primary contig and alternate contig. The primary contig indicates the haplotype, and the alternate contig is composed of short haplotype-specific contigs (haplotigs). Quast v 5.2.0 was used to assess the genome assembly statistics (Gurevich et al. 2013). Gene set completeness was assessed using the Benchmark of Universal Single-Copy Orthologs (BUSCOs) v 5.2.2, with ‘-l insecta_odb10 (2020-09-10)’ (Manni et al. 2021). k-mer-based validation was performed using Merqury v1.3 and Meryl db (k=21) (Rhie et al. 2020) for the assembly constructed with HiFi reads. Contigs were assigned to taxonomic classifications using BlobTools v 1.1(Laetsch and Blaxter 2017) with the BLAST (BLASTN) program in the National Center for Biotechnology Information BLAST software package (v 2.14.0) against the NCBI non-redundant nucleotide sequence database (nt), prepared using ncbi-blast-dbs v 10.5.0 (https://github.com/yeban/ncbi-blast-dbs, accessed on May 18, 2023). minimap2 v 2.26-r1175(Li 2018) was used to obtain mapping coverage.

Dot plots were visualized using the dotPlotly program with ‘-s -t -m 10000 -q 25000 -l’ (https://github.com/tpoorten/dotPlotly, accessed on July 17, 2023) to compare the genomic differences between the US and Japanese strains. The 1 Mb or longer contigs were extracted using SeqKit v 2.0.0, aligned by a nucmer with default settings, and delta files were filtered using a delta-filter with ‘-q -r’ options in MUMmer4 v4.0.0rc (Marçais et al. 2018).

### Gene prediction and functional annotation

Gene prediction was performed using the BRAKER2 and BRAKER3 databases (Brůna et al. 2021; Gabriel et al.). Public RNA sequencing (RNA-Seq) data (morula: DRR138914, DRR138915, DRR138916, and DRR138917; male adult: SRR1864697) and OrthoDB protein sets of Arthropoda (Kuznetsov et al. 2023) (https://bioinf.uni-greifswald.de/bioinf/partitioned_odb11/, accessed on June 17, 2023) were used as extrinsic evidence data to perform the BRAKER pipeline. The RNA-Seq reads were processed using fastp v 0.23.2 with default settings (Chen 2023) and mapped to the *de novo* assembled genome using HISAT2 v 2.2.1(Kim et al. 2019). The generated SAM files were converted to BAM files using SAMtools v 1.12 (Danecek et al. 2021) and the BAM files were imported together when the BRAKER pipeline was run. Repetitive assembly sequences were soft-masked using RepeatModeler v 2.0.3 and RepeatMasker v 4.0.6 in Extensive de novo TE Annotator v 2.1.3 (Ou et al. 2019). The soft-masked genome assembly was inputted into the BRAKER pipeline. GFFRead v0.12.1 (Pertea and Pertea 2020) was used to convert the transcript or protein sequences from the GTF file created by BRAKER. Protein sequences were assessed using BUSCO (protein mode) with ‘-l insecta_odb10’ (2020-09-10).

The functional annotation workflow of Fanflow4Insects (Bono et al. 2022a) was modified and used for protein sequences, as described below. The protein sequences were aligned against reference datasets of *Homo sapiens*, *Mus musculus*, *Caenorhabditis elegans*, *Drosophila melanogaster*, *C. floridanum* (US strain), and UniProtKB/Swiss-Prot using GGSEARCH v 36.3.8g in the FASTA package (https://fasta.bioch.virginia.edu/, accessed on July 19, 2023). Protein sequences were also searched using HMMSCAN in HMMER v 3.3.2. Pfam v 35.0 was used as the protein domain database for HMMSCAN (Mistry et al. 2021). Enrichment analysis was performed using Metascape software (https://metascape.org/, accessed on July 19, 2023) with default settings (Zhou et al. 2019).

## Results and Discussion

### *De novo* genome assembly of Japanese *C. floridanum* and the quality assessment

This study generated HiFi reads and subreads using Japanese strains (Cflo-J). HiFi reads were assembled, and the primary contig comprised 149 contigs with an N50 length of 17.9 Mb and a total assembly size of 552,713,363 bp (50.8-fold coverage; Table 1 and Table S1). Hifiasm also produced alternate contigs (1,162 contigs, Table S1). The genome assembly generated by subreads comprised 996 contigs with an N50 length of 4.6 Mb and a total assembly size of 534,638,157 bp (244.7-fold coverage, Table 1 and Table S1). Cflo_2.0 (NCBI RefSeq assembly GCF_000648655.2) was used as the genome assembly of US strain (Cflo-US). The assembly quality between the Japanese strain (Cflo-J) and the US strain (Cflo-US) was evaluated. The N50 value of Cflo-J assemblies was much higher than that of Cflo-US (1.2 M). However, the GC content of the assemblies was similar. The BUSCO completeness of Cflo-J was also comparable to that of Cflo-US; however, the fragmentation score was lower for the Cflo-J assemblies (Table 1). The N50 values and number of contigs were the best for the HiFi assembly (Table 1 and Figure 1A). We confirmed the differences in genomic structures among all the assemblies using dot-plot visualization (Figures 1B and C). Linear plots were obtained, confirming the similarity of the genomic structures across the entire assembly length (Figure 1B and C). The matching coordinate data for the dot-plot comparison are available in Files S1 and S2.

**Table 1.**
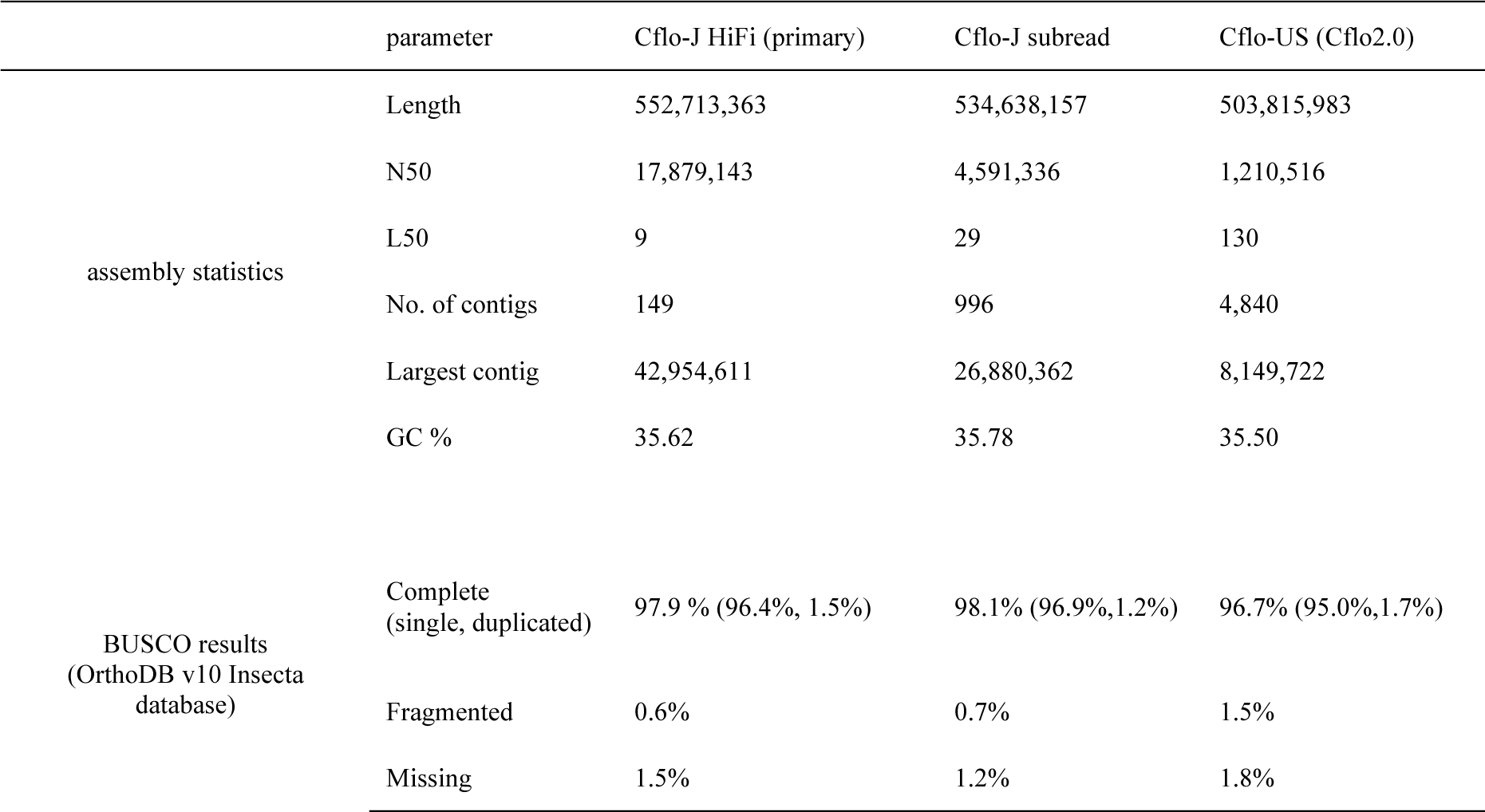
Comparison of assembly statistics with the current reference genomes.

**Figure 1.**
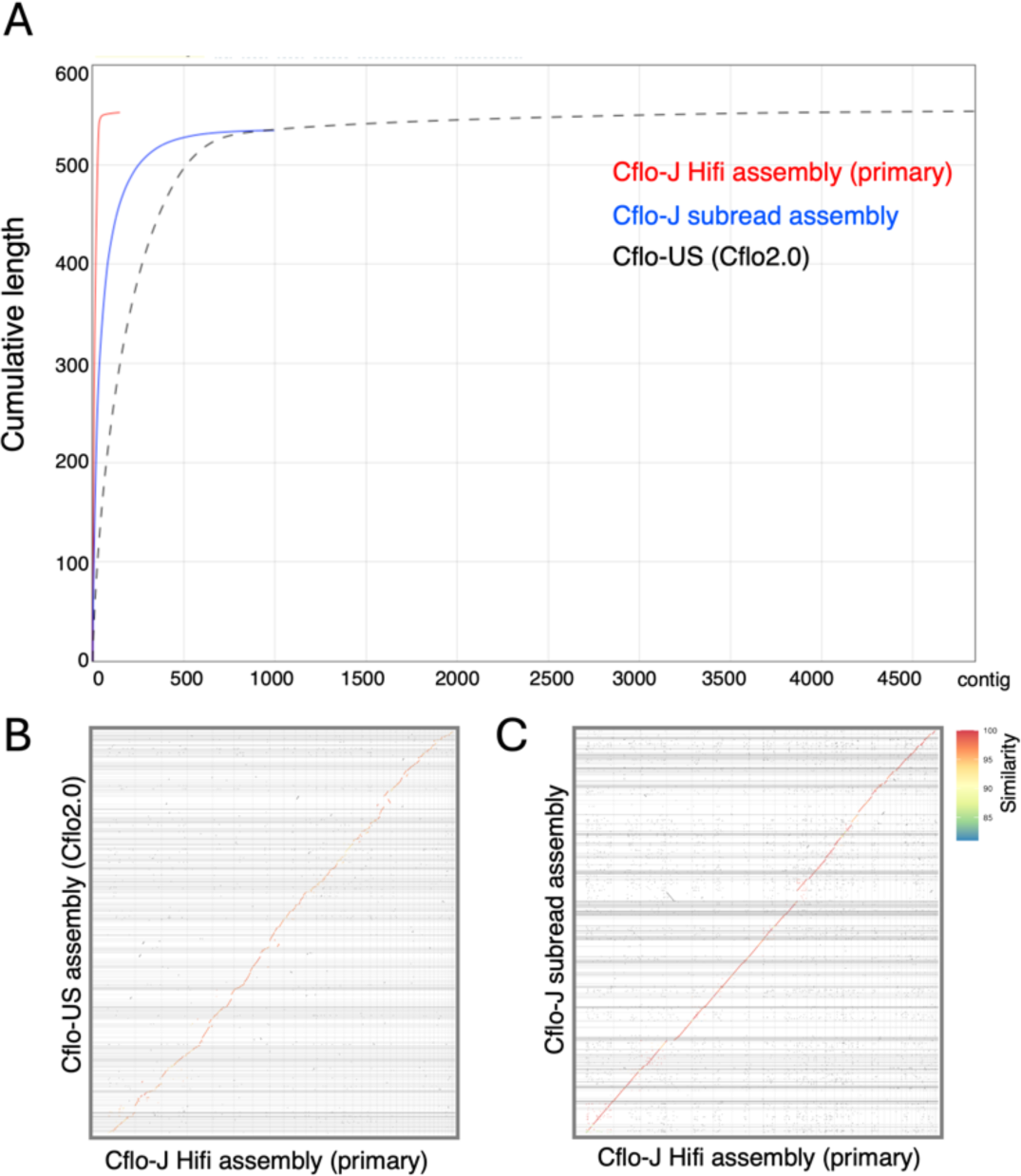
Contiguity and sequence similarity of each assembly. (A) Each color shows the cumulative base pair length and number of contigs for each assembly. The assemblies generated from HiFi reads contain the fewest contigs. (B, C) Dot-plot between the Japanese *C. floridanum* strain (Cflo-J) and US *C. floridanum* (Cflo-US) high-fidelity (HiFi) assemblies and between Cflo-J HiFi and Cflo-J subread assemblies. The vertical axis indicates Cflo-US or Cflo-J subread assemblies. The horizontal axis indicates Cflo-J HiFi assembly. The average identity percentages (per query) are displayed in different colors.

We then evaluated the HiFi assemblies of Cflo-J, which had the highest contiguity. K-mer-based evaluation showed high base accuracy and completeness of primary contigs (a quality value (QV) = 68.08 and completeness = 97.6%, Table S2). The Merqury spectrum plot showed a single k-mer peak in the primary contigs, indicating extremely low heterozygosity (Figure S1A, red line). Because *C. floridanum* males possess a haploid genome, the single k-mer peak of the primary contig was reasonable. We also confirmed the presence of reads included only in the alternate contigs (Figure S1A, blue line), suggesting that the genome assembly constructed in this study may contain heterozygous regions.

Taxonomic assignments of the contigs were performed to reveal the taxonomic origin of the contigs. The primary and alternate contigs were assigned to Arthropoda, and the remaining contigs were either no-hit or chordata (Figures S1B and C). The mapping rate of no-hit was remarkably low (primary contig, 0.27%; alternate contig, 0.51%), and that of chordata in alternate contigs was also extremely low (0.05%). The assembly of Cflo-US also includes Arthropoda and no-hit contigs as shown in Cflo-J (ENA https://www.ebi.ac.uk/ena/browser/view/GCA_000648655.2, accessed on September 20, 2023). In conclusion, an ideal haploid representation was observed in the primary contigs, and no contig contamination from non-arthropods was observed.

### Gene annotation differences between Cflo-J and Cflo-US

As repetitive sequences may affect gene prediction accuracy (Hoff et al. 2019), these sequences in the HiFi primary contig were investigated using RepeatModeler and RepeatMasker. The repeat sequence was 285.4 Mb, corresponding to 51.65% of the HiFi primary contig, and “Unclassified” accounted for a substantial percentage of the total (34.97%) (Table S3). Gene prediction was performed using several RNA-Seq data and protein dataset combinations. The BRAKER3 using RNA-Seq data and the Arthropoda protein dataset showed the best BUSCO completeness (96.4%, Table 2). The numbers of predicted genes and transcripts are listed in Table 2. BRAKER2 predicted more genes compared to BRAKER3 (Table 2). The gene model with the highest BUSCO completeness (96.4%) (“best gene model”) predicted 10,786 protein-coding genes and 13,886 transcripts. The annotated GFF files are shown in File S3. The best gene model was annotated using Fanflow4Insects (Bono et al. 2022b). Table 3 shows the results of the searches using Fanflow4Insects for each reference dataset. Of the 10786 gene models, 10585 were annotated using GGSEARCH or HMMSCAN. The full annotation results for GGSEARCH and HMMSCAN are presented in Table S4.

**Table 2.** BUSCO analysis for the protein set predicted by BRAKER.

| Table 2 BUSCO analysis for the protein set predicted by BRAKER |  |  |  |  |
| --- | --- | --- | --- | --- |
|  | BRAKER3 with RNA seq<br>+ Arthropoda protein | BRAKER2 with RNA<br>seq | BRAKER3 with<br>Arthropoda protein | BRAKER2 with<br>Arthropoda protein |
| Complete<br>(single, duplicated) | 96.4 % (79.5%, 16.9%) | 94.0% (73.9%, 20.1%) | 93.1% (78.8%, 14.3%) | 92.5% (82.8%, 9.7%) |
| Fragmented | 0.5% | 2.4% | 0.7% | 1.8% |
| Missing | 3.1% | 3.6% | 6.2% | 5.7% |
| No. of genes | 10786 | 18531 | 9913 | 22781 |
| No. of transcripts | 13886 | 19986 | 12370 | 24487 |

**Table 3.**
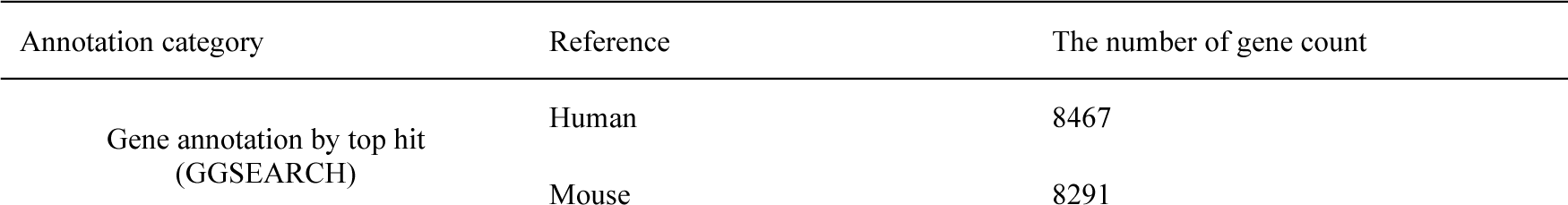
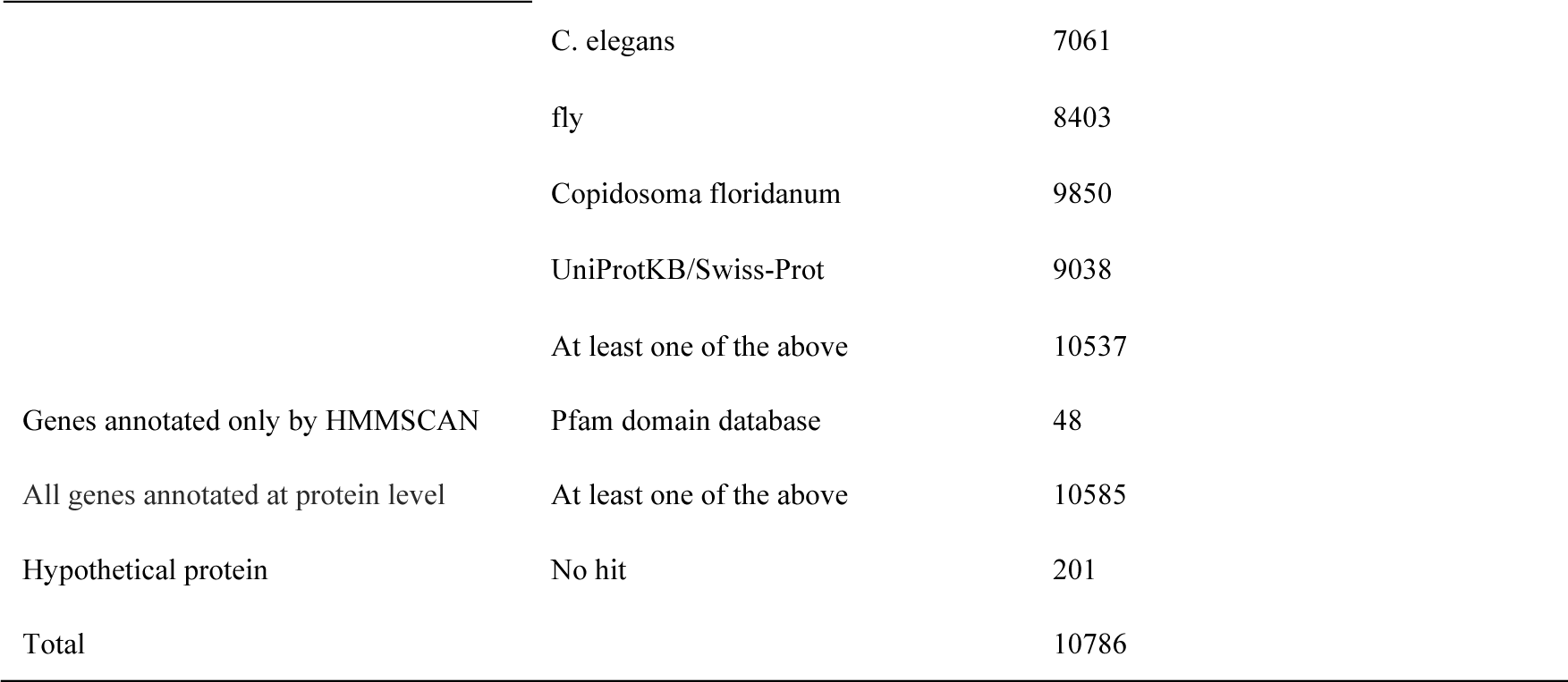
Summary of functional annotation results.

| Table 3 Summary of functional annotation results |  |  |
| --- | --- | --- |
| Annotation category | Reference | The number of gene count |
| Gene annotation by top hit<br>(GGSEARCH) | Human | 8467 |
|  | Mouse | 8291 |
|  | C. elegans | 7061 |
|  | fly | 8403 |
|  | Copidosoma floridanum | 9850 |
|  | UniProtKB/Swiss-Prot | 9038 |
|  | At least one of the above | 10537 |
| Genes annotated only by HMMSCAN | Pfam domain database | 48 |
| All genes annotated at protein level | At least one of the above | 10585 |
| Hypothetical protein | No hit | 201 |
| Total |  | 10786 |

We evaluated whether annotation differences existed between the Cflo-US assembly and Cflo-J HiFi assembly. We identified 149 transcripts in 115 gene loci with no corresponding proteins to Cflo-US and with corresponding proteins to human, mouse, *C.elegans*, and fly (Table S5). The human gene IDs corresponding to 149 transcripts were used for enrichment analysis (Figure 2A). “Metabolism of carbohydrates” was the most significantly enriched Gene Ontology (GO) term. In addition, GO terms for amino acid metabolism and translation were enriched.

**Figure 2.**
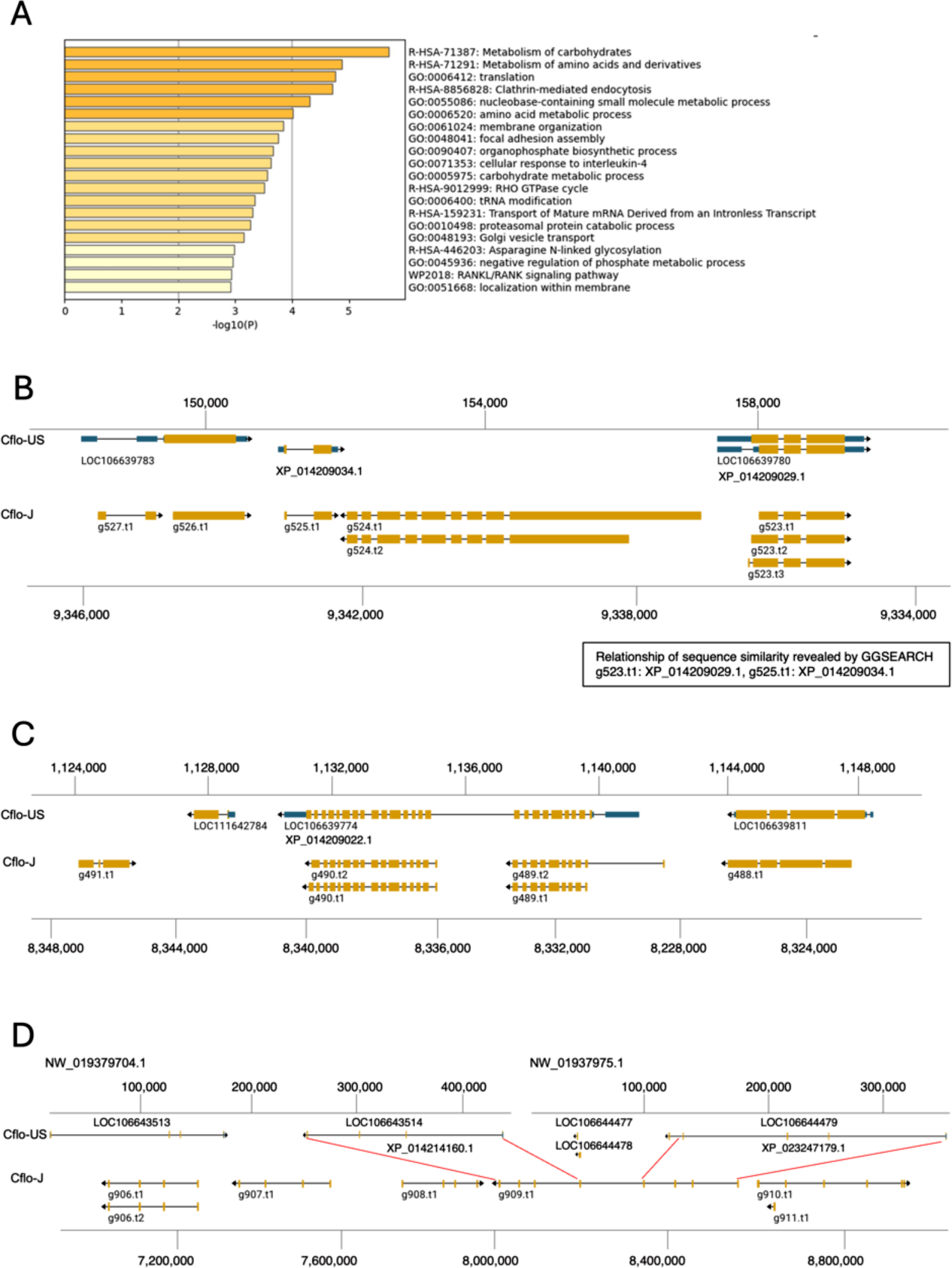
Features of genes with no corresponding proteins to Cflo-US. (A) The results of enrichment analysis (Metascape) of 149 transcripts with no corresponding proteins to Cflo-US and with corresponding proteins to human, mouse, *C. elegans*, and fly. Bars are colored differently according to the value of - log10 (P-value). (B-D) Representative gene model illustrating the genome annotation differences. Yellow and blue boxes indicate the ORF and UTR, respectively. Red lines indicate the region with similar exon structures.

Excluding gene loci with at least one isoform corresponding to Cflo-US genes, the number of genes as mentioned above was limited from 115 to 100. We investigated why these 100 genes did not hit the Cflo-US gene model. To verify whether corresponding regions of the 100 genes are absent in the Cflo-US, TBLASTN of 100 genes was performed for the genome of Cflo-US. We manually examined hit regions of TBLASTN using NCBI Genome Data viewer. TBLASTN results showed that all 100 genes hit in the Cflo-US genome. Thus, the difference in genome annotation between Cflo-US and Cflo-J is not due to differences in the extent of the sequenced genomic regions. Among 100 genes, 21 genes had no corresponding gene models in Cflo-US assembly, while 79 genes had corresponding gene models in Cflo-US (Table S6). The gene model g524 was representative example without corresponding gene model of Cflo-US (Figure 2B). On both sides of g524 are g523 and g525. Amino acid sequences of g523 and g525 have sequence similarity to XP_014209029.1 and XP_014209034.1 in Cflo-US genome, respectively (Table S4). Although these genomic position and sequence similarity indicate that the genomic position around g524 is conserved in both genome assembly, the gene model corresponding to g524 was absent in Cflo-US.

Among 79 genes with corresponding gene model in Cflo-US, TBLASTN results were divided into two groups: genes that hit the single gene model or genes that hit more than two gene model (Table S6). In addition, differences of the amino acid length between Cflo-J and Cflo-US were also revealed by the TBLASTN (Table S6). g490 was representative example as the gene that hit the single gene model of Cflo-US. TBLASTN showed that the amino acid sequence of g490.t1 had similarity to XP_014209022.1 of Cflo-US gene model (Table S6). The exon structures of XP_014209022.1 were different from g490 of Cflo-J (Figure 2C). Multiple sequence alignment showed that the amino acid length of XP_014209022.1 of Cflo-US was extremely longer at N-terminal region than the corresponding protein of other species (Figure S2). When BLASTP was adopted, g490.t1 showed high sequence similarity to XP_014209022.1 (E-value: 0.0), despite the gene structures were largely different. This indicates that GGSEARCH adopted in this study can detect the genes with extremely different structure of genes. g909 was the representative example as the gene that hit several gene model of Cflo-US (Table S6). TBLASTN showed that g909 had the similarity XP_023247179.1 and XP_014214160.1, and these two genes of Cflo-US were distributed in different scaffolds (Figure 2D). Multiple sequence alignment indicated that XP_023247179.1 and XP_014214160.1 correspond to the first and second half of the amino acid sequence of g909.t1, respectively (Figure S3). The example of g909 suggested that long-read sequencing contributed to the accuracy of genome annotation. TBLASTN also detected the genes that hit more than two gene model on the same scaffold (Table S6, e.g. g1061.t1).

In this study, we found genes with extremely different gene annotations between Cflo-US and Cflo-J. This success is attributed to the method used in this study. GGSEARCH, which was adopted in this study, searches for sequence similarity with the subject within 80-120% of the query length. This contributed to discovery of protein structure differences between Cflo-J and Cflo-US. In addition, highly contiguous genome assembly obtained by long-read sequencing suggested the fragmentation of gene model in Cflo-US genome. All the differences in genome annotation revealed in this study are noteworthy as the discoveries of genomic differences between Cflo-J and Cflo-US.

### Common features and differences of *vasa* between Cflo-J and Cflo-US

Ohno et al. (2019) showed that the amino acid sequence of *vasa* differed between Cflo-J and Cflo-US. The amino acid sequences of Cflo-US (XP_014219851.1) and Cflo-J *vasa* (g8014.t1), including the amino acid sequence (BBI30140.1) determined by Ohno et al. (2019), were compared. We confirmed the differences in amino acid sequences at the N-terminal region between Cflo-US *vasa* (XP_014219851.1) and Cflo-J *vasa* (g8014.t1), as previously reported^13^ (Figure 3A, red arrows). Within the Japanese strain (BBI30140.1 vs. g8014.t1), the amino acid sequences were almost identical (Figure 3A).

**Figure 3.**
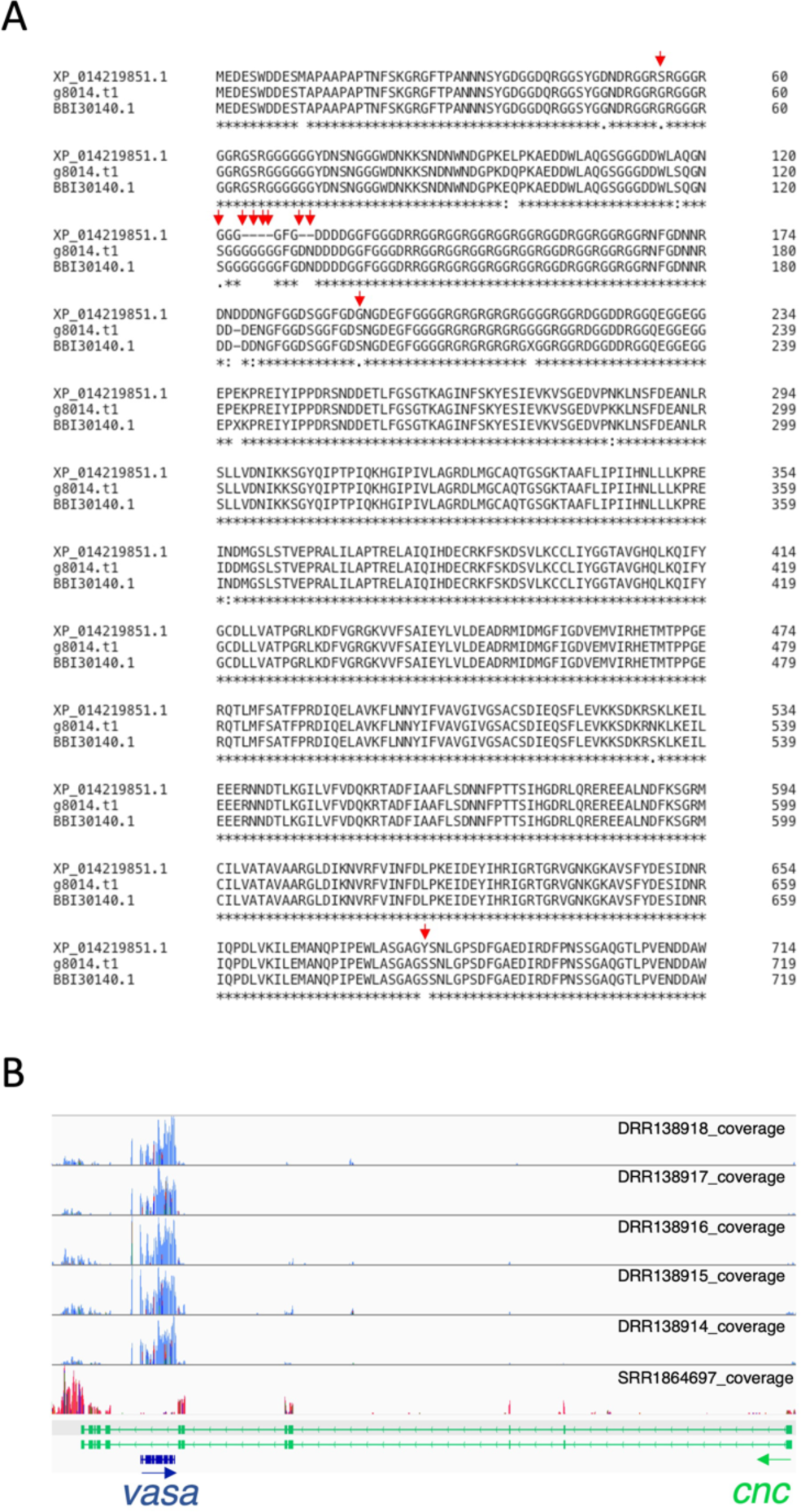
Amino acid sequences genomic position and of *vasa* gene. Comparison of gene annotation between the Cflo-US and Cflo-J HiFi assemblies. (A) Comparison of amino acid sequences of the *vasa* gene in the US (XP_014219851.1) and Japanese strains (g8014.t1 and BBI30140.1). g8014.t1 is the transcript ID of *vasa* predicted in Cflo-J assembly, and BBI30140.1 is the protein ID of *vasa* in the Japanese lineage reported by Ohno et al. (2019). Red arrows indicate amino acids that differ between the US and Japanese strains, as Ohno et al. 2019 revealed. (B) Genomic position of *vasa* with RNA-Seq read coverages. The CDS regions are shown at the bottom, including the *vasa* and *cap’n’collar* (*cnc*) exon structures. Blue and green arrows indicate the direction of transcription.

We also investigated the genomic position of *vasa* (Figure 3B). *vasa* was located in an intron between the seventh and eighth exons of *cap’n’collar* (*cnc*), and the directions of transcription were opposite (Figure 2C, blue and green arrows). This positional relationship was also confirmed in Cflo-US (NCBI Genome Browser: LOC106647823, Location: scaffold6|NW_019379589.1:2,626,762 – 2,630,760.). Generally, overlapping genes such as *vasa* and *cnc* are negatively selected evolutionarily, whereas overlapping genes are sometimes retained in a species-specific manner (Craig et al. 1997). In addition, overlapping genes are likely to be co-expressed. Furthermore, the *vasa* genomic positions in other insects were preliminarily examined (Table S7), and the positional relationship between *vasa* and *cnc* may be conserved in Hymenoptera. Genomic positional relationship of *vasa* and *cnc* may be related to the oogenesis or embryogenesis in hymenopteran insects. *cnc* is related to the nuclei arrangement of oocyte in *Drosophila melanogaster* (Guichet et al. 2001), suggesting its involvement in egg development in *C. floridanum*. The evolution to control the caste development by *vasa* in *Copidosoma* may have required a conserved positional relationship between *vasa* and *cnc* in the Hymenoptera. However, further verifications for relationship between *vasa* and *cnc* are needed.

## Conclusion

We obtained a highly contiguous genome assembly of Japanese *C. floridanum*. We showed that long-read sequencing and functional annotation method adopted in this study can highlight the differences of genome annotation between Cflo-J and Cflo-US. These regions may be due to the influence of the annotation and assembly accuracy, but in any case, they are remarkable genes that reflect differences among the strains. Parasitic wasps can control the physiological state of host insects, and their use for pest control is expected (Soldà et al. 2008). Comparison of the genome sequences of Cflo-US and Cflo-J will contribute to the identification of detailed genomic regions relevant to the ecological and physiological characteristics of *C. flolidanum*, leading to a detail understanding of its parasitic ability against pests.

## Data availability

All sequencing data (assembled sequences and raw sequence reads) were deposited in the DDBJ under the umbrella BioProject accession number PRJDB16642. Genome assembly from primary contigs was deposited in the DDBJ under the accession numbers BTPR01000001‒BTPR01000149. Alternate contigs were deposited in DDBJ under the accession numbers BTBT001162. Raw sequence reads were deposited in the DDBJ under accession number DRA017002 (PacBio HiFi reads) and DRA017125 (PacBio Subreads). Gene annotation and functional annotation of protein-coding genes are available at figshare (https://doi.org/10.6084/m9.figshare.24145182.v5). Supplementary Files are available at figshare (https://doi.org/10.6084/m9.figshare.24145182.v5). Scripts of genome assembly, quality check, and gene annotation were also available (https://doi.org/10.6084/m9.figshare.24145182.v5)

## Acknowledgments

We would like to thank all laboratory members at Hiroshima University for their valuable comments. Computations were performed on the computers at the Hiroshima University Genome Editing Innovation Center. Computations were partially performed on the NIG supercomputer at ROIS National Institute of Genetics.

## Authors’ contributions

Conceptualization: H.T. and H.B.; methodology: Kouhei T. and H.B.; software: Kouhei T. Formal analysis, data curation, and visualization: Kouhei T., Keita T., and T.S. Validation: Kouhei T., and H.B. Investigation: Kouhei T., and H.B. Resources: Kouhei T., H.B., and H.T. Writing—original draft: Kouhei T. Writing—review and editing: T.S., M.K, Keita T., K.O., H.T., and H.B. Supervision: H.T., and H.B. Project administration: H.B. Funding acquisition: H.T., and H.B.

## Conflict of interest

The authors declare no conflict of interest.

## Funding

This study was supported by the Center of Innovation for Bio-Digital Transformation (BioDX), an open innovation platform for industry-academia co-creation of JST (COI-NEXT, grant number JPMJPF2010 to H.T. and H.B.), and JSPS KAKENHI (grant number 23K17418 to H.T. and H.B.).

## Notes

https://doi.org/10.6084/m9.figshare.24145182.v5

### Competing Interest Statement

The authors have declared no competing interest.

### Summary of Updates

The results for differences between U.S. and Japanese strains were added significantly.

## References

Ardui S, Ameur A, Vermeesch JR, Hestand MS. 2018. Single molecule real-time (SMRT) sequencing comes of age: applications and utilities for medical diagnostics. Nucleic Acids Res. 46(5):2159–2168. doi:10.1093/nar/gky066.

Bono H, Sakamoto T, Kasukawa T, Tabunoki H. 2022a. Systematic Functional Annotation Workflow for Insects. Insects. 13(7):586. doi:10.3390/insects13070586.

Bono H, Sakamoto T, Kasukawa T, Tabunoki H. 2022b. Systematic Functional Annotation Workflow for Insects. Insects. 13(7):586. doi:10.3390/insects13070586.

Brůna T, Hoff KJ, Lomsadze A, Stanke M, Borodovsky M. 2021. BRAKER2: automatic eukaryotic genome annotation with GeneMark-EP+ and AUGUSTUS supported by a protein database. NAR Genom Bioinform. 3(1). doi:10.1093/nargab/lqaa108.

Chen S. 2023. Ultrafast one-pass FASTQ data preprocessing, quality control, and deduplication using fastp. iMeta. 2(2). doi:10.1002/imt2.107.

Cheng H, Concepcion GT, Feng X, Zhang H, Li H. 2021. Haplotype-resolved de novo assembly using phased assembly graphs with hifiasm. Nat Methods. 18(2):170–175. doi:10.1038/s41592-020-01056-5.

Cook JM. 1993. Sex determination in the Hymenoptera: a review of models and evidence.

Craig SF, Slobodkin LB, Wray GA, Biermann CH. 1997. The ‘paradox’ of polyembryony: A review of the cases and a hypothesis for its evolution. Evol Ecol. 11(2):127–143. doi:10.1023/A:1018443714917.

Cruz YP. 1981. A sterile defender morph in a polyembryonic hymenopterous parasite. Nature. 294(5840):446–447. doi:10.1038/294446a0.

Danecek P, Bonfield JK, Liddle J, Marshall J, Ohan V, Pollard MO, Whitwham A, Keane T, McCarthy SA, Davies RM, et al. 2021. Twelve years of SAMtools and BCFtools. Gigascience. 10(2). doi:10.1093/gigascience/giab008.

Donnell DM, Corley LS, Chen G, Strand MR. 2004. Caste determination in a polyembryonic wasp involves inheritance of germ cells. Proc Natl Acad Sci U S A. 101(27):10095–10100. doi:10.1073/pnas.0403625101.

Eid J, Fehr A, Gray J, Luong K, Lyle J, Otto G, Peluso P, Rank D, Baybayan P, Bettman B, et al. 2009. Real-Time DNA Sequencing from Single Polymerase Molecules. Science (1979). 323(5910):133–138. doi:10.1126/science.1162986.

Gabriel L, Brůna T, Hoff KJ, Ebel M, Lomsadze A, Borodovsky M, Stanke M. BRAKER3: Fully Automated Genome Annotation Using RNA-Seq and Protein Evidence with GeneMark-ETP, AUGUSTUS and TSEBRA. doi:10.1101/2023.06.10.544449. https://doi.org/10.1101/2023.06.10.544449.

Giron D, Harvey JA, Johnson JA, Strand MR. 2007. Male soldier caste larvae are non-aggressive in the polyembryonic wasp *Copidosoma floridanum*. Biol Lett. 3(4):431–434. doi:10.1098/rsbl.2007.0199.

Grbic M, Rivers D, Strand MR. 1997. Caste formation in the polyembryonic wasp Copidosoma floridanum (Hymenoptera: Encyrtidae): in vivo and in vitro analysis. J Insect Physiol. 43(6):553–565. doi:10.1016/S0022-1910(97)00004-8.

Guerrieri E, Noyes J. 2005. Revision of the European species of Copidosoma Ratzeburg (Hymenoptera: Encyrtidae), parasitoids of caterpillars (Lepidoptera). Syst Entomol. 30(1):97–174. doi:10.1111/j.1365-3113.2005.00271.x.

Guichet A, Peri F, Roth S. 2001. Stable anterior anchoring of the oocyte nucleus is required to establish dorsoventral polarity of the drosophila egg. Dev Biol. 237(1):93–106. doi:10.1006/dbio.2001.0354.

Gurevich A, Saveliev V, Vyahhi N, Tesler G. 2013. QUAST: Quality assessment tool for genome assemblies. Bioinformatics. 29(8):1072–1075. doi:10.1093/bioinformatics/btt086.

Hay B, Yeh Jan L, Nung Jan Y. 1988. A Protein Component of Drosophila Polar Granules Is Encoded by vasa and Has Extensive Sequence Similarity to ATP-Dependent Helicases.

Hoff KJ, Lomsadze A, Borodovsky M, Stanke M. 2019. Whole-Genome Annotation with BRAKER. Methods Mol Biol. 1962:65–95. doi:10.1007/978-1-4939-9173-0_5.

Hotaling S, Sproul JS, Heckenhauer J, Powell A, Larracuente AM, Pauls SU, Kelley JL, Frandsen PB. 2021. Long Reads Are Revolutionizing 20 Years of Insect Genome Sequencing. Genome Biol Evol. 13(8). doi:10.1093/gbe/evab138.

Hotaling S, Wilcox ER, Heckenhauer J, Stewart RJ, Frandsen PB. 2023. Highly accurate long reads are crucial for realizing the potential of biodiversity genomics. BMC Genomics. 24(1):117. doi:10.1186/s12864-023-09193-9.

i5K Consortium. 2013. The i5K Initiative: advancing arthropod genomics for knowledge, human health, agriculture, and the environment. J Hered. 104(5):595–600. doi:10.1093/jhered/est050.

Iwabuchi K. 2019. Polyembryonic Insects. Singapore: Springer Singapore.

Kawahara AY, Storer CG, Markee A, Heckenhauer J, Powell A, Plotkin D, Hotaling S, Cleland TP, Dikow RB, Dikow T, et al. 2022. Long-read HiFi sequencing correctly assembles repetitive heavy fibroin silk genes in new moth and caddisfly genomes. GigaByte. 2022:1–14. doi:10.46471/gigabyte.64.

Kim D, Paggi JM, Park C, Bennett C, Salzberg SL. 2019. Graph-based genome alignment and genotyping with HISAT2 and HISAT-genotype. Nat Biotechnol. 37(8):907–915. doi:10.1038/s41587-019-0201-4.

Kolmogorov M, Yuan J, Lin Y, Pevzner PA. 2019. Assembly of long, error-prone reads using repeat graphs. Nat Biotechnol. 37(5):540–546. doi:10.1038/s41587-019-0072-8.

Kuznetsov D, Tegenfeldt F, Manni M, Seppey M, Berkeley M, Kriventseva EV, Zdobnov EM. 2023. OrthoDB v11: annotation of orthologs in the widest sampling of organismal diversity. Nucleic Acids Res. 51(D1): D445–D451. doi:10.1093/nar/gkac998.

Laetsch DR, Blaxter ML. 2017. BlobTools: Interrogation of genome assemblies. F1000Res. 6:1287. doi:10.12688/f1000research.12232.1.

Lasko PF, Ashburner M. 1988. The product of the Drosophila gene vasa is very similar to eukaryotic initiation factor-4A. Oxford University Press.

Li H. 2018. Minimap2: pairwise alignment for nucleotide sequences. Bioinformatics. 34(18):3094– 3100. doi:10.1093/bioinformatics/bty191.

Manni M, Berkeley MR, Seppey M, Simão FA, Zdobnov EM. 2021. BUSCO Update: Novel and Streamlined Workflows along with Broader and Deeper Phylogenetic Coverage for Scoring of Eukaryotic, Prokaryotic, and Viral Genomes. Mol Biol Evol. 38(10):4647–4654. doi:10.1093/molbev/msab199.

Marçais G, Delcher AL, Phillippy AM, Coston R, Salzberg SL, Zimin A. 2018. MUMmer4: A fast and versatile genome alignment system. PLoS Comput Biol. 14(1): e1005944. doi: 10.1371/journal.pcbi.1005944.

Mistry J, Chuguransky S, Williams L, Qureshi M, Salazar GA, Sonnhammer ELL, Tosatto SCE, Paladin L, Raj S, Richardson LJ, et al. 2021. Pfam: The protein families database in 2021. Nucleic Acids Res. 49(D1): D412–D419. doi:10.1093/nar/gkaa913.

Ode PJ, Strand MR. 1995. Progeny and Sex Allocation Decisions of the Polyembryonic Wasp Copidosoma floridanum.

Ohno H, Sakamoto T, Okochi R, Nishiko M, Sasaki S, Bono H, Tabunoki H, Iwabuchi K. 2019. Apoptosis-mediated vasa down-regulation controls developmental transformation in Japanese Copidosoma floridanum female soldiers. Dev Biol. 456(2):226–233. doi: 10.1016/j.ydbio.2019.09.005.

Ou S, Su W, Liao Y, Chougule K, Agda JRA, Hellinga AJ, Lugo CSB, Elliott TA, Ware D, Peterson T, et al. 2019. Benchmarking transposable element annotation methods for creation of a streamlined, comprehensive pipeline. Genome Biol. 20(1):275. doi:10.1186/s13059-019-1905-y.

Pertea G, Pertea M. 2020. GFF Utilities: GffRead and GffCompare. F1000Res. 9:304. doi:10.12688/f1000research.23297.2.

Pollard MO, Gurdasani D, Mentzer AJ, Porter T, Sandhu MS. 2018. Long reads: their purpose and place. Hum Mol Genet. 27(R2): R234–R241. doi:10.1093/hmg/ddy177.

Raz E. 2000. The function and regulation of vasa-like genes in germ-cell development. Genome Biol. 1(3): REVIEWS1017. doi:10.1186/gb-2000-1-3-reviews1017.

Rhie A, Walenz BP, Koren S, Phillippy AM. 2020. Merqury: reference-free quality, completeness, and phasing assessment for genome assemblies. Genome Biol. 21(1):245. doi:10.1186/s13059-020-02134-9.

Rhoads A, Au KF. 2015. PacBio Sequencing and Its Applications. Genomics Proteomics Bioinformatics. 13(5):278–289. doi: 10.1016/j.gpb.2015.08.002.

Sakamoto T, Nishiko M, Bono H, Nakazato T, Yoshimura J, Tabunoki H, Iwabuchi K. 2020. Analysis of molecular mechanism for acceleration of polyembryony using gene functional annotation pipeline in Copidosoma floridanum. BMC Genomics. 21(1):152. doi:10.1186/s12864-020-6559-3. https://bmcgenomics.biomedcentral.com/articles/10.1186/s12864-020-6559-3.

Schupbach T, Wieschaus E. 1986. Maternal-effect mutations altering the anterior-posterior pattern of the Drosophila embryo. Roux’s Archives of Developmental Biology. 195(5):302–317. doi:10.1007/BF00376063.

Segoli M, Harari AR, Rosenheim JA, Bouskila A, Keasar T. 2010. The evolution of polyembryony in parasitoid wasps. J Evol Biol. 23(9):1807–1819. doi:10.1111/j.1420-9101.2010.02049.x.

Smith MS, Shirley A, Strand MR. 2017. Copidosoma floridanum (Hymenoptera: Encyrtidae) Rapidly Alters Production of Soldier Embryos in Response to Competition. Ann Entomol Soc Am. 110(5):501–505. doi:10.1093/aesa/sax056.

Soldà G, Suyama M, Pelucchi P, Boi S, Guffanti A, Rizzi E, Bork P, Tenchini ML, Ciccarelli FD. 2008. Non-random retention of protein-coding overlapping genes in Metazoa. BMC Genomics. 9. doi:10.1186/1471-2164-9-174.

Strand MR. 2003. Polyembryony. Carde R., Resch V., editors. San Diego: Academic Press.

Szathmáry E, Smith JM. 1995. The major evolutionary transitions. Nature. 374(6519):227–232. doi:10.1038/374227a0.

Uka D, Takahashi-Nakaguchi A, Yoshimura J, Iwabuchi K. 2013. Male soldiers are functional in the Japanese strain of a polyembryonic wasp. Sci Rep. 3(1):2312. doi:10.1038/srep02312.

Vladimir Zhurov, Tomislav Terzin, Miodrag Grbic. 2004. Early blastomere determines embryo proliferation and caste fate in a polyembryonic wasp. Nature. 432(7018):761–764. doi:10.1038/nature03030.

Whyte BA. 2021. The weird eusociality of polyembryonic parasites. Biol Lett. 17(4). doi:10.1098/rsbl.2021.0026.

Yamamoto D, Henderson R, Corley LS, Iwabuchi K. 2007. Intrinsic, inter-specific competition between egg, egg–larval, and larval parasitoids of plusiine loopers. Ecol Entomol. 32(2):221–228. doi:10.1111/j.1365-2311.2006.00857.x.

Zhou Y, Zhou B, Pache L, Chang M, Khodabakhshi AH, Tanaseichuk O, Benner C, Chanda SK. 2019. Metascape provides a biologist-oriented resource for the analysis of systems-level datasets. Nat Commun. 10(1):1523. doi:10.1038/s41467-019-09234-6. http://www.nature.com/articles/s41467-019-09234-6.

